# Advanced adaptive strategies in an ancestral body plan: insights from a 510-Ma-old leptomitid sponge

**DOI:** 10.1101/2025.05.30.657026

**Authors:** Cui Luo, Yanjie Hong, Zhixin Sun, Haijing Sun, Swee-Cheng Lim, Tianyu Wang, Lei Zhang, Fangchen Zhao

**Affiliations:** Key Laboratory of Palaeobiology and Petroleum Stratigraphy, Chinese Academy of Sciences, Nanjing 210008, People’s Republic of China; University of Chinese Academy of Sciences (UCAS), Beijing 100049, People’s Republic of China; Lee Kong Chian Natural History Museum, National University of Singapore, 2 Conservatory Drive, Singapore 117337, Singapore; Institute of Sedimentary Geology, Chengdu University of Technology, Chengdu 610059

**Keywords:** Ascospongiae, protomonaxonid, modularity, functional morphology, siliciclastic, Guanshan Biota

## Abstract

Sponges have thrived in diverse environmental conditions since the early Cambrian till today. However, little is known about how their adaptive capability and strategies have been shaped throughout evolutionary history. Here, we explore this question based on a new leptomitid sponge fossil from the Cambrian Stage 4. The family Leptomitidae was an abundant sponge group inhabiting Cambrian soft substrates but significantly declined thereafter. The new species exhibits a sophisticated set of morphological characteristics adaptive to a shallow siliciclastic environment, which are unprecedented among leptomitids. These include (1) a robust body wall woven by spirally twisted monaxonic spicules; (2) a thick stub-like root tuft for anchoring; (3) spicules radiating out from the sponge body to prevent clogging and sinking; and (4) the inferred capability to close the osculum against unfavourable stimuli. Nevertheless, the new fossil species maintains a leptomitid body plan which lacks modularity and morphological plasticity, the two common and critical attributes in extant sponges to enhance flexibility and resilience in changing environmental conditions. This juxtaposition of evolutionary innovation and structural conservatism offers a compelling case for further exploration of the evolutionary mechanisms that shaped early sponge lineages.

## INTRODUCTION

Sponges are an ancient animal phylum deeply rooted in the Neoproterozoic^1,2^ and have been successful throughout the Phanerozoic (e.g., Kiessling et al.^3^). Today, sponges are the second most diversified non-bilaterian phylum after Cnidaria, with more than 9400 valid species distributed from freshwater lakes and shallow marine reefs to abysses^4,5^. In fact, their adaptability to a broad range of environments has been achieved since the earliest undisputable sponge fossil representatives. Sponge spicules have been discovered from shallow water carbonates to slope and basinal cherts in the Fortunian (539–529 Ma)^6–10^. Archaeocyaths, an extinct group of hypercalcified sponges, were flourishing in the early Cambrian reefs during ca. 529–509 Ma^11^. Meanwhile, from Cambrian Stage 3 to Drumian (521–500.5 Ma), articulated skeletal frames of spicular sponges are one of the major fossil types in the shale Lagerstätten distributed from shoreface-offshore to basinal settings^12–18^. Although the adaptive tactics and mechanisms have been intensively investigated in living sponges^19–23^, much less is known regarding how these features evolved over time. Have these strategies remained largely unchanged throughout the over-500-Myr evolutionary history, or have they been progressively shaped and refined over this long period?

The family Leptomitidae is an ancient sponge group which was prevalent and abundant in the Cambrian siliciclastic settings^17,25,26,33,50,51,74^ but significantly declined after the Cambrian and then the Ordovician^24^. Little is known about the causes of their thriving and disappearance^24^, although both the fluctuating environmental factors and the biology of the organisms *per se* must have played roles. For instance, all the leptomitids known so far possess a pointed lower end^25^, indicating that they were sediment stickers adapted to a Proterozoic-type substrate, which is characterized by a well-developed microbial mat covering, low levels of bioturbation, and low water content^27^. It is conceivable that the increasing dominance of the Phanerozoic- to the Proterozoic-type soft substrate during the Cambrian-Ordovician interval posed a challenge for sponges retaining such an ancient anchoring strategy^28^.

In this study, we examine the growth mode and functional morphology of a new sponge fossil from the Guanshan Biota, Cambrian Age 4 (514–509 Ma). This fossil, assigned to the family Leptomitidae, exhibits an exceptionally sophisticated skeletal architecture within the family. Through this case study, we aim to demonstrate how complex and effective adaptive morphological variations can emerge within the constraints of a conservative leptomitid body plan.

## RESULTS

### Preservation of the sponge fossils

Most of the investigated sponge fossils were collected from the Wulongqing Formation in the Xiaopengzu section in Luquan County, eastern Yunnan province (Figure 1A), co-occurring with trilobites and hyoliths correlative with the middle part of the Wulongqing Formation and early Cambrian Stage 4 (see Materials and Methods; Figures 1B–1J). The other specimens, donated by amateur palaeontologists and loaned from museums, were also collected from the ambient area.

**Figure 1.**
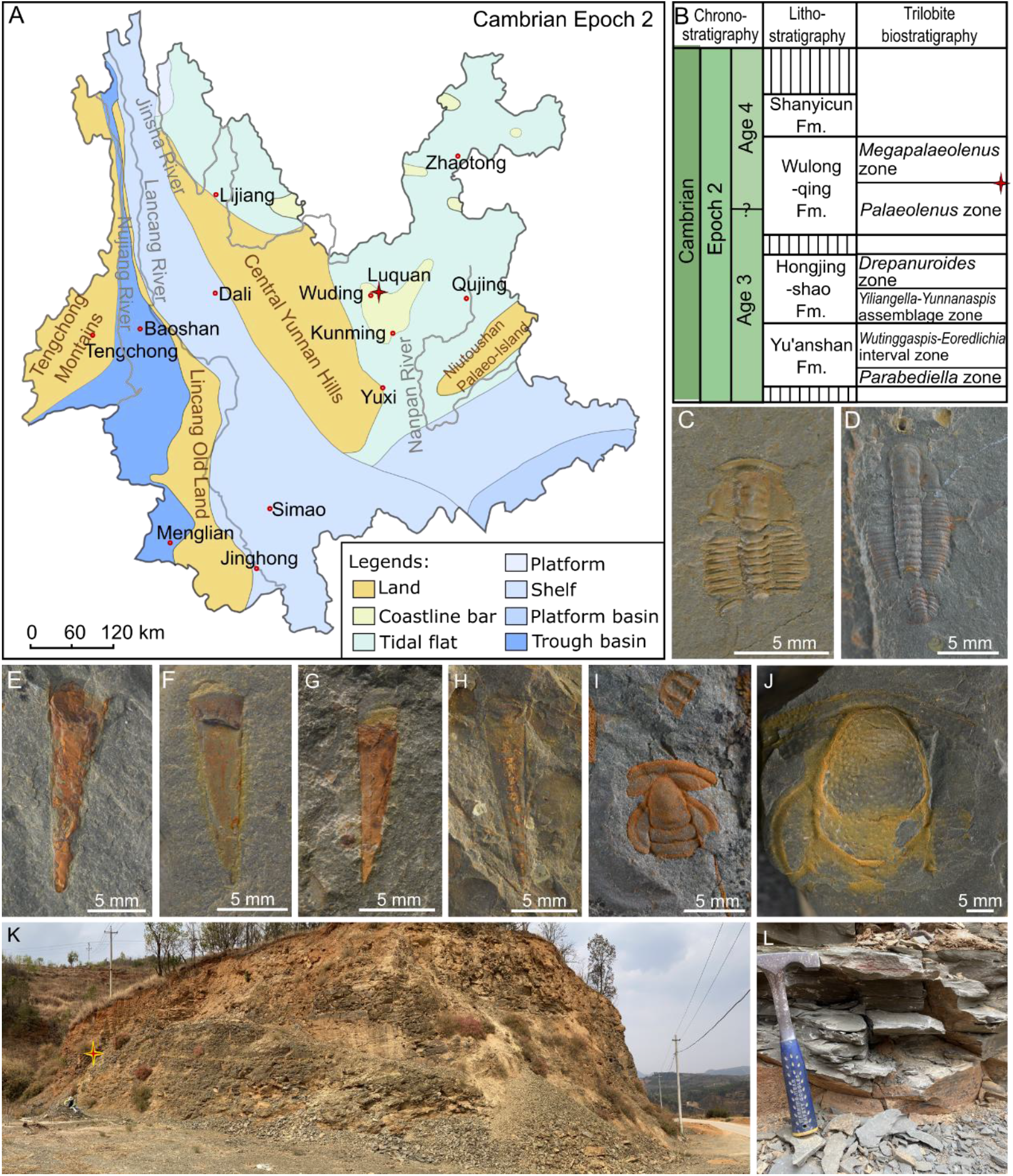
(A) Sedimentary facies and palaeogeography of Yunnan Province in Cambrian Epoch 2, after Yunnan Bureau of Geology and Mineral Resources^29^. The star indicates Luquan County, where the Xiaopengzu section is located. (B) Litho- and biostratigraphy of the Cambrian Series 2 of Wuding City, eastern Yunnan, after Steiner et al.^30^ and Zhu et al.^31^. The star indicates the investigated fossil level. (C–J) Fossils co-occurring with the studied sponges: *Megapalaeolenus deprati* (C, D), “*Linevitus*” *guizhouensis* (E, F), *Doliutheca orientalis* (G, H); *Redlichia yunnanensis* (I), *Breviredlichia granulosa* (J). (K, L) The outcrop of the Xiaopengzu section.

The Wulongqing Formation is dominated by storm-influenced silty and muddy deposits except for the basal conglomerates. It mainly represents an offshore environment in eastern Yunnan^32,33^. The sponge fossils are preserved in a yellowish-greenish silty shale succession (Figures 1K and 1L), and the most fossiliferous levels are associated with enriched skeletal fragments of other animals. Thin sections show that the fossil-bearing host rock is poorly laminated and contains silty patches with diffuse boundaries (Figures 2C and 2D). This structure is comparable to that of the deposits accumulated from the suspension of depositional events^38^. Together, these observations imply an involvement of transportation before the burial of these fossils. The sponge fossils are strongly flattened and often stained by iron oxides (Figures 2A–2D). Spicules have lost their original composition and are mainly preserved as reliefs inseparable from the host rock (Figures 2E–2J). These fossils, therefore, represent the external casts and moulds of the organism.

**Figure 2.**
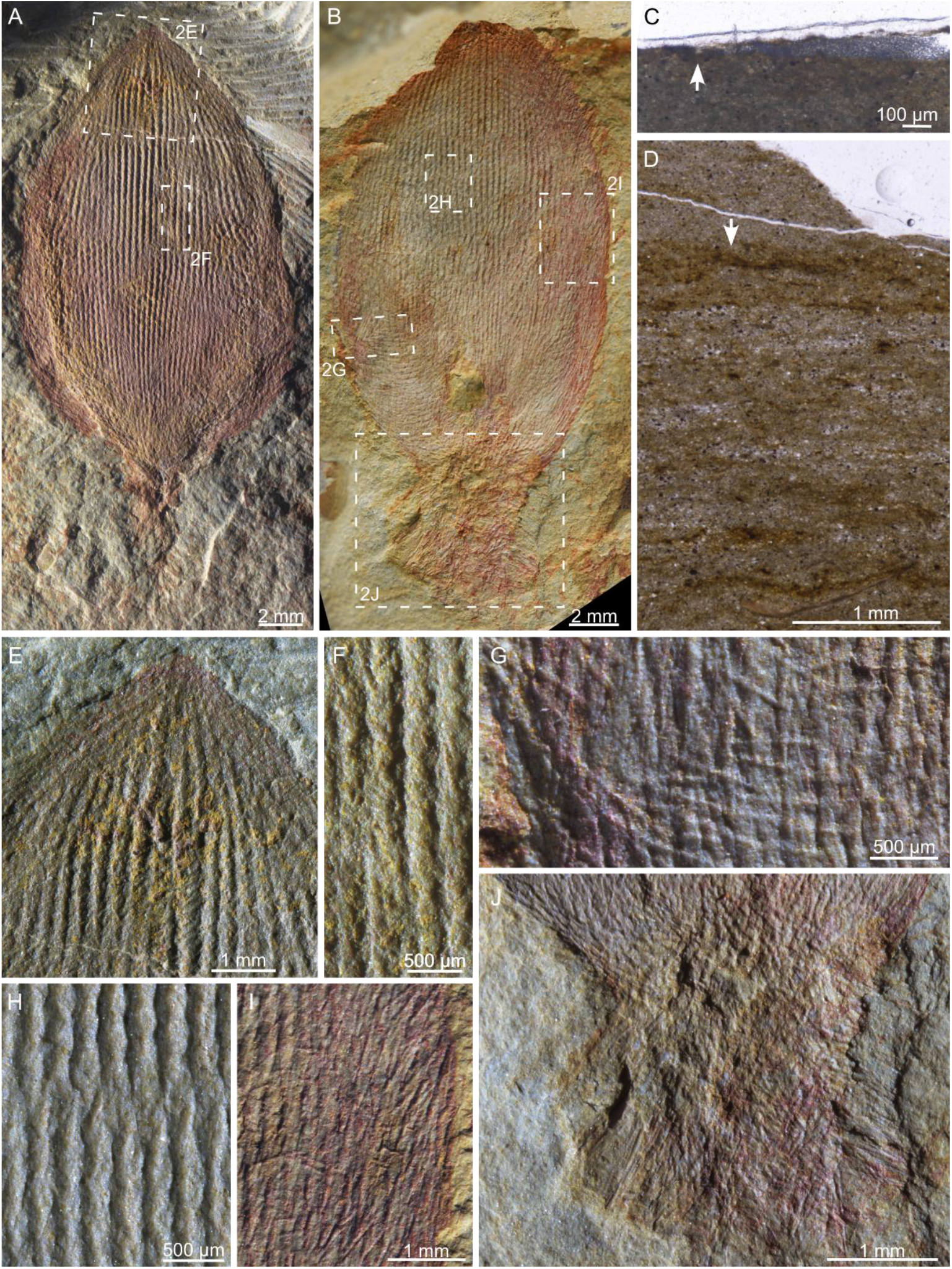
(A, E, F) The holotype of *Lotispongia helicolumna*, FMD02a-01. (B, G, J) A paratype of *L. helicolumna*, YL-2-02. (C, D) Thin sections from the hand specimen xpz230318-32. The arrow in C indicates the cross-section of a bud-like body. The wedge of silty filling below the epoxy on the right part of the image is an artefact during processing. The arrow in D indicates the same layer where the fossil in C occurs.

### Systematic Palaeontology

**Phylum:** Porifera Grant^34^

**Class:** Uncertain

**Order:** ‘‘Protomonaxonida’’ Finks & Rigby^35^ *sensu* Botting et al.^36^

**Family:** Leptomitidae de Laubenfels^37^

**Genus:** *Lotispongia* gen. nov. Luo

Type species: *Lotispongia helicolumna* sp. nov. Luo

#### Etymology

“Loti-”, Latin, refers to the shape of the sponge that looks like a lotus flower bud.

#### Diagnosis

The fossil is a small, non-branching sponge with a lotus-bud-shaped body and a stub-like root tuft. The spongocoel, with its osculum extremely reduced, is surrounded by a thin body wall which is primarily constructed by densely arranged longitudinal spicule bundles. Each spicule bundle is formed by spirally twisted long monaxons. Short horizontal monaxons reinforcing the spicule bundles may or may not be present. Monaxons of intermediate length radiate upwards and outwards from the body wall. Similar monaxons assemble to form the root tuft but radiate downwards.

#### Remarks

The morphology of this sponge conforms with the diagnosis of the family Leptomitidae in being mainly constructed by bundling, long longitudinal spicules supplemented by discrete horizontal elements (Figure 3). However, the lotus-bud-like body, the thick root tuft, and the additional monaxons radiating from the body wall readily separate this genus from other leptomitids^25,35,44,50,51^ (Figure 3). In addition, the extremely reduced osculum and the longitudinal spicule bundles made by spirally twisted monaxons were also unknown in leptomitids. Nevertheless, these unique characteristics can be interpreted as morphological variations within the family. It does not seem necessary to erect a new family for this single genus at this stage.

**Figure 3.**
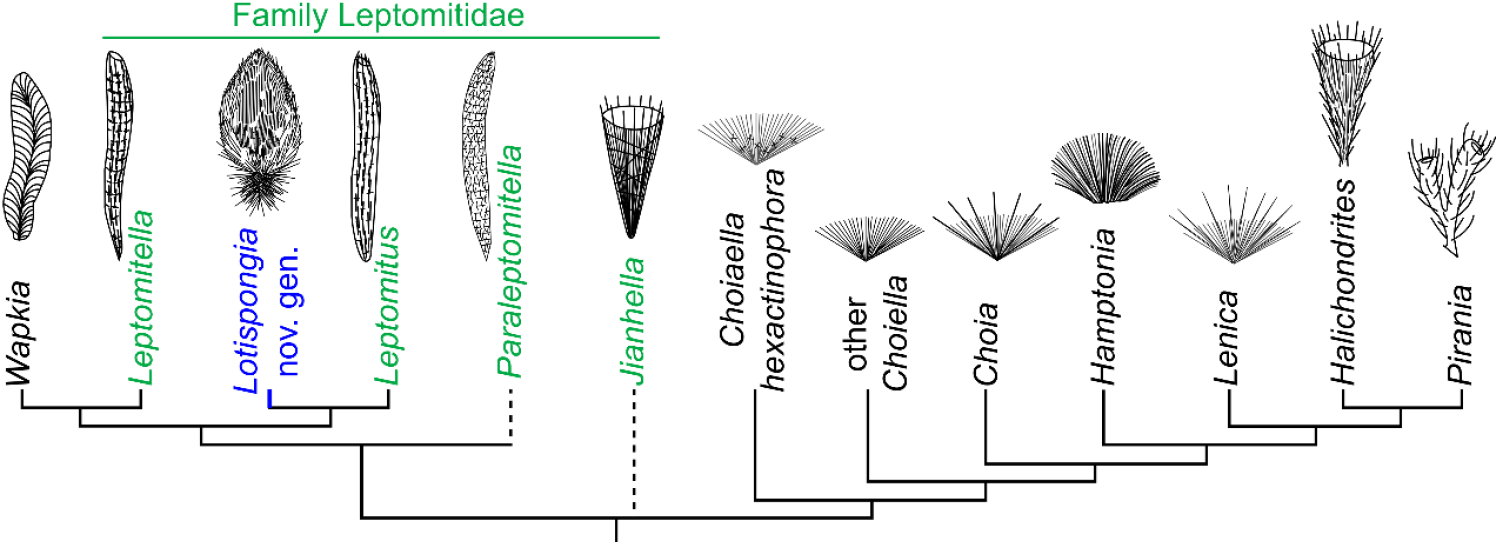
Morphology of major Group One (PG1) protomonaxonid genera and their assumed phylogenetic relationships according to Botting et al.^36^, Wang et al.^50^, and Botting^39^. The sketches of the fossils were adapted or created based on published literature^39,50–52^. *Hyalosinica* has been excluded from this frame according to Yun et al.^53^.

**Figure 4.**
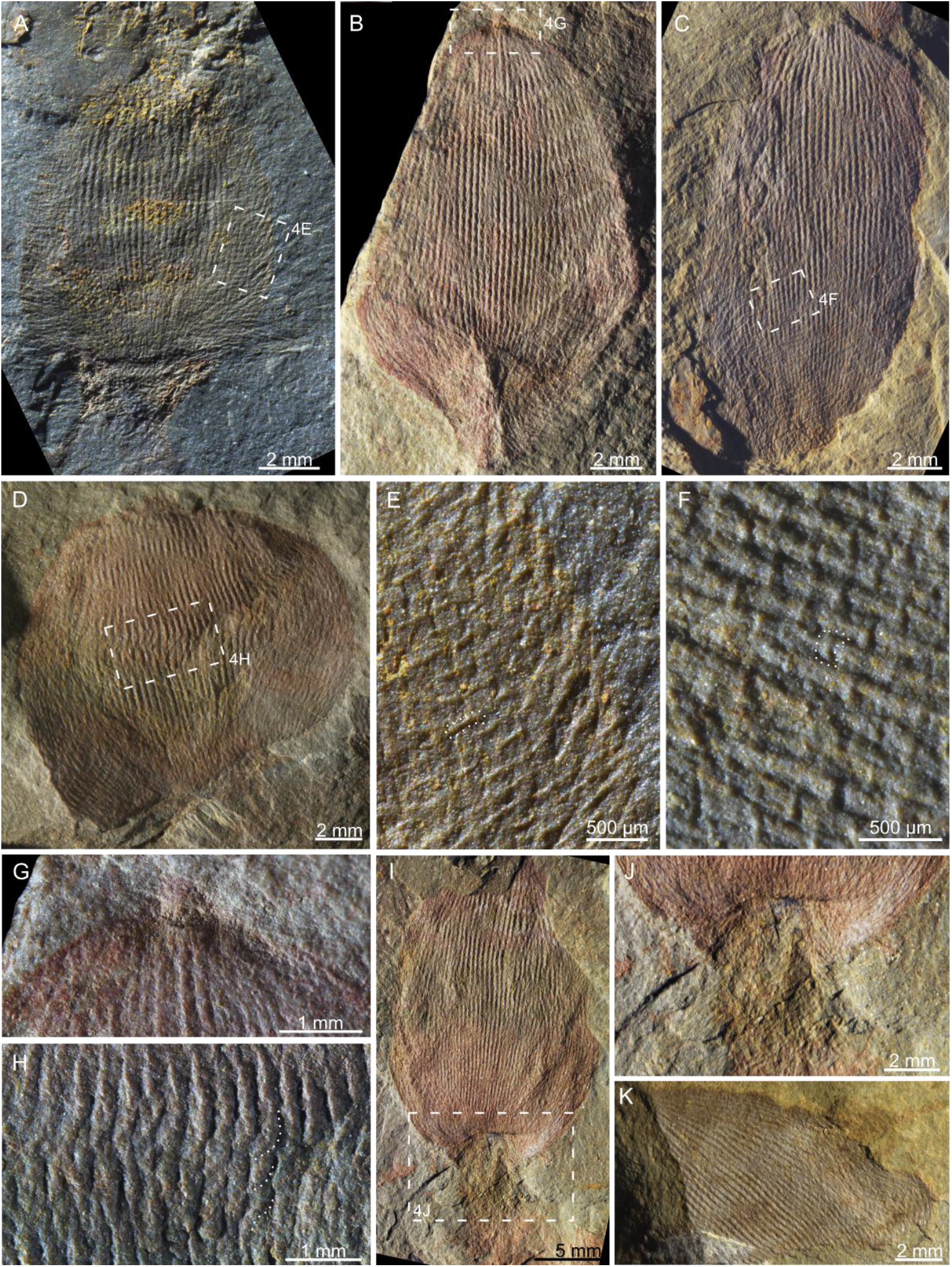
Morphological details in a few paratypes of *Lotispongia helicolumna*, including the short horizontal monaxons (E, F), the fossil top (G), the rope-like longitudinal spicule bundles (H), the connection between the body and the root tuft (I, J), and the distortion of the sponge body (D, K). The dotted lines in E, F, and H highlight examples of the respective structures that are intended to be illustrated. Article number: (A, E) xpz20230318-26-01; (B, G) FMD02b-02; (C, F) KMDLa-02; (D, H) FMD01-02; (I, J) FMD03-02; (K) YL-1-03.

With such a readily determined family-level taxonomic solution, this new genus’s order- and class-level taxonomy is more controversial. The order Protomonaxonida Finks & Rigby^35^, to which the family Leptomitidae previously belonged, was a poorly defined taxon even in terms of morphological classification. Botting et al.^36^ separated this order into two groups: Group One (hereafter PG1) is characterized by large longitudinal spicules with an unresolved affinity, while Group Two (hereafter PG2) is composed of complex tracts of small monaxons and considered to be early demosponges. A new class, Ascospongiae, was recently proposed as a formal replacement of the Protomonaxonida and encompasses PG1^39^.

Based on skeletal architecture, we concur that PG1 and PG2 can be recognized as distinct groups. However, the proposed establishment of a new class-level taxon for PG1 warrants a more careful and critical evaluation.

First, the skeletal architecture alone cannot be regarded as an adequate argument to erect a new class for PG1, as based on architecture, the PG1 fossils can be and have been interpreted as extinct groups belonging to either class Demospongiae^35^ or class Hexactinellida^40^. The most decisive argument for erecting the new class is the alleged capability of ascosponges to produce distinct spicules which “may be open-based and partly hollow in derived groups, with prominent organic sheath externally.”^39^ However, these spicule characteristics observed from different fossils may need verification to exclude their alternative origins from primary biological characteristics. For instance, the open-based spicule in *Pirania* (Figures 5A and 5B in Botting & Muir^41^) can also be a fragmented chancelloriid sclerite buried with the specimen and does not belong to the sponge (Figure S1). Organic matter surrounding fossil spicules can have several other origins instead of representing primary sheaths enveloping the spicules. For instance, it can derive from remaining organic matter from the cellular soft tissue, diagenetic polymerization of organic matter in the skeleton^42^, or even carbonaceous material migrated into the fossil-associated pores from the host rock^43^. It has been observed that both macro- and microscopic mineral skeletal fossils can demineralize and be replaced by carbon^45,46^.

**Figure 5.**
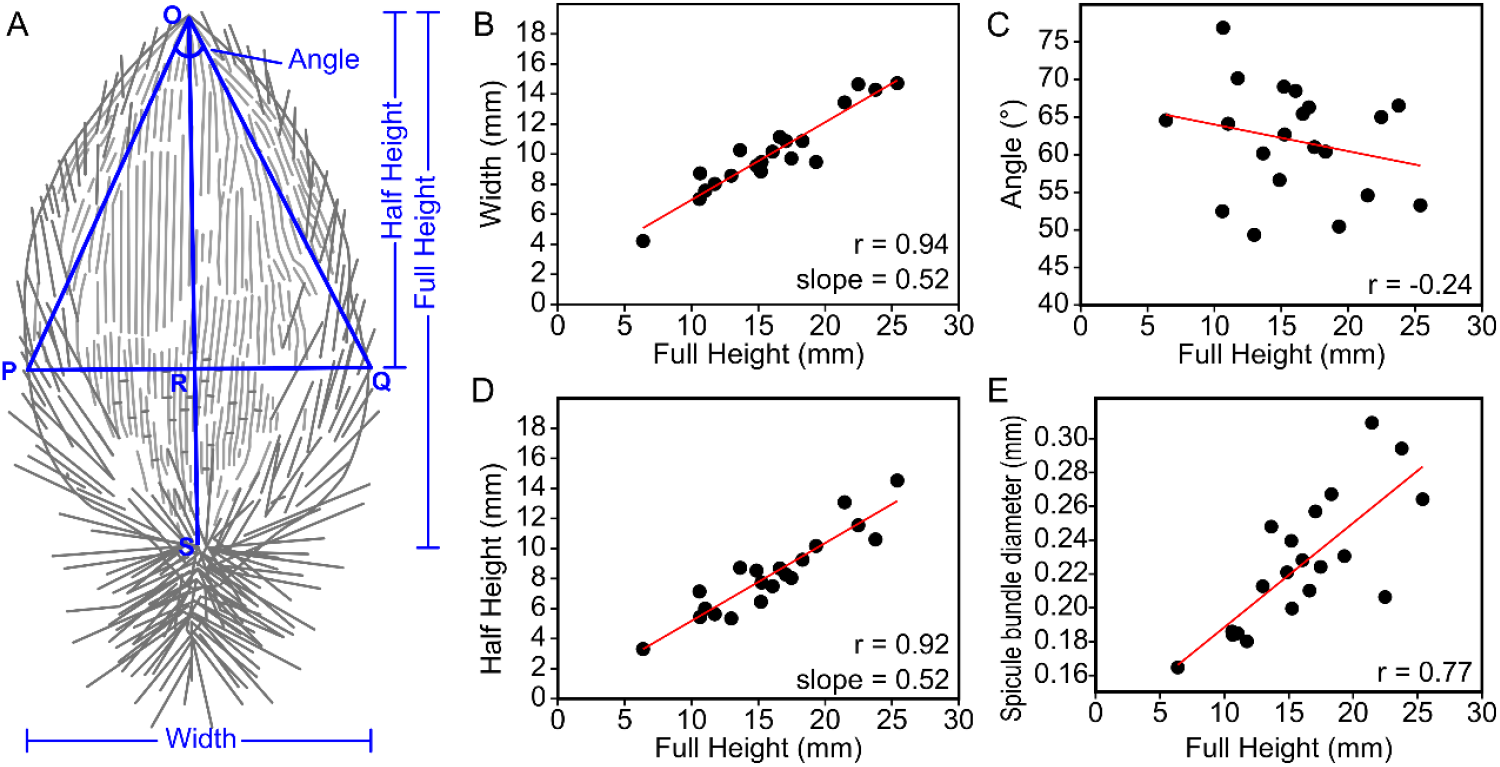
Morphometric analysis of *Lotispongia helicolumna* based on 20 specimens in which all the variables are measurable. (A) shows the meaning of the variables “Full Height”, “Half Height”, “Width”, and “Angle”. The letters O, P, Q, R, and S are used to name the nodes in the geometric shape. (B–E) Linear correlations between Full Height and other parameters.

Moreover, although not written in the diagnosis, some spicules of ascosponges were interpreted as bimetallic^39,47^, inferred from the different degrees of mineral dissolution in fossil materials^47^. However, studies of extant spicules showed that the core of a spicule can be dissolved faster than the peripheral^48^, and this cavity can be rapidly filled by secondary mineral precipitation^49^. Diagenetic processes often involve generations of dissolution and precipitation. It’s plausible that the secondarily precipitated minerals in early diagenesis can be further altered during deeper burial. Interpretations of atypical biomineralization characteristics require careful scrutiny when derived from diagenetically modified material. Our observation of a well-preserved spicule fossil from the Cambrian Stage 3 black shale also shows that the concentric layers were preserved in different silica phases that show varied solubility (Figure S2).

Given the concerns outlined above, we suggest that a more comprehensive evaluation of the ascosponge concept is needed. However, such an investigation is beyond the scope of this study. Therefore, for the time being, we solve the class- and order-level taxonomy of *Lotispongia* following the way of Botting & Brayard (2019)^24^.

**Species:** *Lotispongia helicolumna* sp. nov. Luo (Figures 2, 4)

#### Etymology

“helic-”, Latin, means spiral; “columna”, Latin, means column. Together, “helicolumna” refers to the rope-like spicule bundles in the body wall.

#### Specimens

Holotype, FMD02-01a, b (Figures 2A, 2E, 2F); Paratypes, 24 specimens listed in Table S1; Other materials include 55 specimens, also listed in Table S1.

#### Locality and horizon

Luquan County, Kunming, Yunnan Province; Cambrian Stage 4.

#### Diagnosis

As for the genus.

#### Description

The bud-like body of the sponge is 6.4–25.4 mm tall (mean = 16.0 ± 4.8 mm, n = 23) and 4.2– 14.7 mm wide (mean = 9.9 ± 2.6 mm, n = 25). The largest exposure of the root tuft is 13.2 mm long and 7.6 mm wide, attaching to a 25.4 mm × 14.7 mm bud-like body (FMD08).

No unambiguous open osculum was observed in these fossils (Figures 2A, 2B, 2E, 4A– 4D, 4G). Opening-like structures were only found at the fossil top of FMD02b-02 and FMD14- 01 among the 80 investigated specimens, and both cases can be alternatively interpreted as products of early diagenetic distortion and/or excavation breakage (Figure 4G). Nevertheless, *L. helicolumna* probably possesses a large spongocoel. This is inferred from the considerable number of specimens whose body was softly deformed (Figures 4D and 4K), indicating that the bud-like body was composed of a thin wall surrounding a hollow space. In comparison, the root tuft is probably a solid assembly of spicules, for it appears to be less flattened than the body, was never deformed, and often penetrates the sediments (Figure 2J). The contrast of stiffness is especially conspicuous at the junction of the body and the root tuft (Figures 4A, 4B, 4I, 4J).

The body wall is basically composed of long longitudinal spicule bundles, which are 117– 309 μm (mean = 210 ± 43 μm, n = 77) wide and consist of spirally twisted spicules (Figures 2E, 2F, 2H, 4H). In some places, the spicules in the rope-like spicule bundles disentangle and then join with spicules from neighbouring spicule bundles (Figure 2H). Short horizontal spicules of 41–108 μm thick (mean = 73 ± 19 μm, n = 20) are observed only in a few specimens, especially noticeable at the lower part of the organism (Figures 4E and 4F).

Attached to the body wall are upward and outward radiating monaxons. The spicules, 42– 102 μm wide (mean = 74 ± 19 μm, n = 15) and up to 1.13 mm long, can be abundantly distributed throughout the whole body (Figures 2B, 4A) or be preserved only at the base and edge of the fossil (Figures 2A, 2E, 4B, 4C, 4G). This indicates that these spicules may have been less tightly articulated with the body wall and can be shed off or washed away before burial. The spicules in the root tuft are similar in shape and size to these radiating spicules but extend outwards and downwards (Figures 2A and 2B, 4A).

#### Remarks

The rope-like spicule bundles composed of spirally twisted spicules are uncommon among fossil sponges but known from *Kiwetinokia*, which was mainly described from the Cambrian Stage 4–Drumian of North America^54–56^. However, *Kiwetinokia* was assigned to reticulosans or hexactinellids, and their rope-like spicule bundles were interpreted as anchoring spicules. The occurrence of similar spicule bundles in the parenchymal skeleton of *L. helicolumna* is probably an independent innovation of this leptomitid sponge.

### Morphometric analyses

The regular organization of leptomitid skeletons has been observed for a long time^36^. However, most leptomitid fossils are not preserved with a complete outline due to their centimetric to decimetric length^25^. For this reason, the speculated skeletal regularity in leptomitids has never been studied using quantitative methods. The small size and robust architecture of *L. helicolumna* (discussed in detail below), for the first time, allows such an analysis.

Five parameters were considered to characterize the morphology of *L. helicolumna* (Figure 5A): (1) the height of the bud-like body (Full Height = length of OS), (2) the largest width of the body (Width = length of PQ), (3) the distance from the top of the sponge to the widest part (Half Height = length of OR), (4) the vertex angle of the triangle that connects the sponge top and the two points of the sponge’s outline where the body reaches the widest part (Angle = ∠O), and (5) the average diameter of the rope-like spicule bundles.

Based on data from 20 almost completely preserved specimens (Table S2), Width against Full Height and Half Height against Full Height both show conspicuous linear correlations (Figs. 5B, 5D). This means that OR and PQ in Figure 5A change almost proportionally with OS and both maintain a length ratio of around 0.5 relative to the latter (Figs. 5B, 5D). These correlations also indicate that ∠O approaches a fixed value as the sponge size increases. The measured data show that Angle does not change relative to Full Height and is mainly distributed in the range of 50°–70° (Figure 5C). The diameter of the rope-like spicule bundles is also linearly correlated with the body size (Figure 5E).

Although all measured values have undoubtedly been overprinted by burial deformation, the results nonetheless reveal a clear pattern: *L. helicolumna* maintained a constant geometry throughout its growth.

## DISCUSSION

### Effective adaptive characteristics in *Lotispongia helicolumna*

Unlike other sponges whose skeletal frame disarticulates soon after the organism dies, the skeleton of the studied specimens of *L. helicolumna* is preserved almost entirely after transportation and soft deformation, without evidence of tearing and disintegration, as shown in Figures 4D and 4H. The mechanical robustness can be attributed to the tightly twisted longitudinal spicules. More remarkably, each “rope” can disentangle and then weave into neighbouring rope-like spicule bundles (Figure 2H). This organization enables the sponge to maintain the stability of the skeletal frame in agitated water regardless of the presence or absence of supplementary horizontal spicules.

The thick root tuft of *L. helicolumna* suggests that it is a soft-bottom dweller^22^, in accordance with its burial environment. The monaxons radiating outwards from the body are also known as a strategy in living sponges to prevent sediments from clogging lateral inhalant pores^22^. In some specimens, the radiating monaxons at the upper part of the root tuft and the lower part of the body extend para-horizontally (Figures 2J, 4A). This feature can prevent the sessile organism from sinking into the loose sediment^22^.

The most peculiar part of *L. helicolumna* is its lotus-bud-like body, which is composed of a bulged base and a pointed apical end devoid of an observable osculum. The pointed apical end is uncommon in the phylum Porifera but does exist on some occasions. Firstly, some sponges have a stable growth form with a pointed top, such as the extant hexactinellid species *Semperella sjades* Lim & Setiawan^57^ and the fossil hexactinellid *Sanshapentella tentoriformis* Yun et al.^58^. Although its ecological benefits are unclear, this morphology seems to be permanent in *Semperella sjades*. The permanence of the pointed apical end in *Sanshapentella tentoriformis* can also be determined because this structure is supported by specialized tent-like large pentactins^58^. The extant soft-bottom-dwelling demosponge *Craniella quirimure* sometimes also shows a lotus-bud-like shape, but most of them are described as ovoid to spherical^59^.

Secondly, many living demosponges are capable of closing their oscula during unfavourable stimulations^60^, and the closed oscula can appear as cones without obvious openings^22,61,62^(Figure 6A). This seems to be a more probable interpretation for the pointed apical end in *L. helicolumna*. Unlike *Semperella sjades* and *Sanshapentella tentoriformis, L. helicolumna* is a thin-walled sponge with a single large spongocoel. An osculum is a necessity in such an aquiferous system. In contrast, *Semperella sjades* lacks a single large exhalant cavity. Instead, its complex aquiferous system expels the spent water through multiple oscula across the atrial areas^57^. *Sanshapentella tentoriformis* is also a thick-walled sponge, branching, and does not have an obvious major osculum. It is plausible that *Sanshapentella tentoriformis* possessed a complex aquiferous system with multiple small oscula, similar to that of *Semperella sjades*. However, if *L. helicolumna* had a single osculum, the normal size of its osculum could not have been as small as being unrecognizable. For hydrodynamic efficiency, the optimal oscular diameter should be of intermediate size, not approaching zero^63–65^. Taking all these arguments together, we infer that the pointed apical end of *L. helicolumna* represents an osculum closed due to stimuli, instead of a fixed morphology. To our knowledge, this is the first report of osculum contraction in a sponge fossil, despite this behaviour being well known in extant sponges^22,61,62^(Figure 6A).

**Figure 6.**
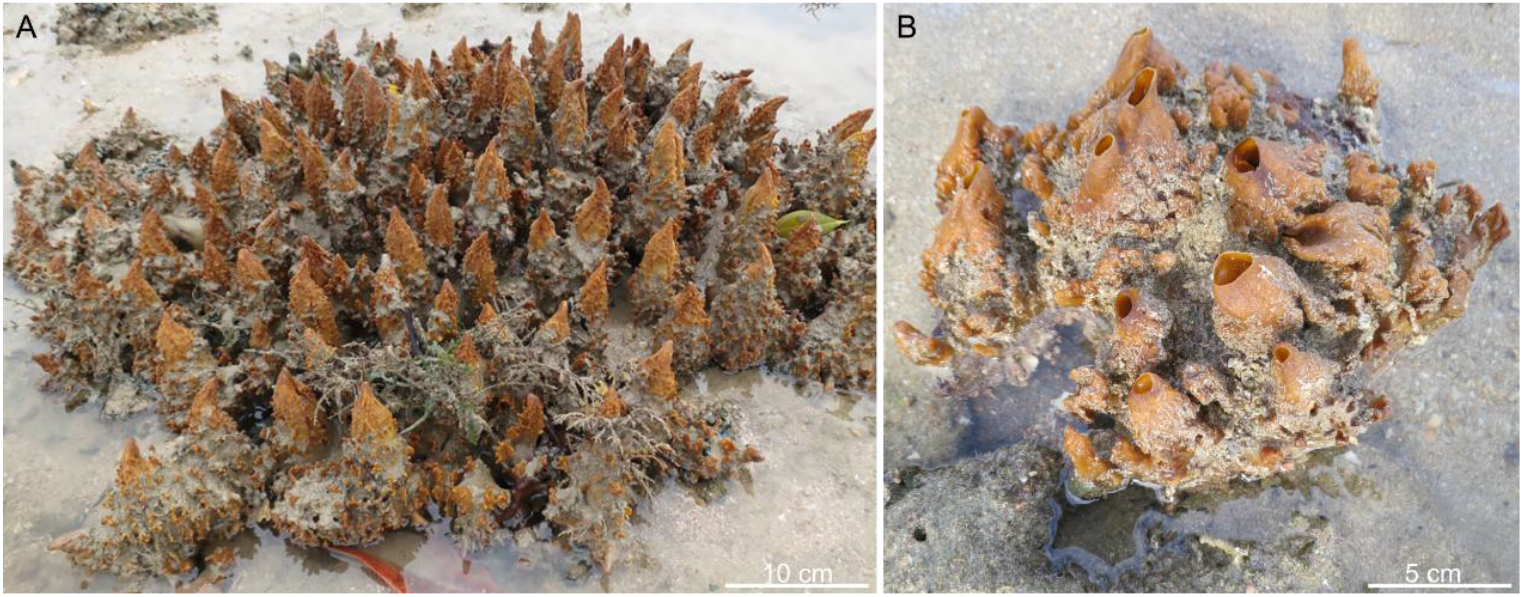
Endopsammic demosponge *Spheciospongia* sp. from Singapore, with the oscula closed (A) and open (B). A is reproduced from Figure 1c in Schönberg & Lim^62^.

### Possible drawbacks of the *Lotispongia helicolumna* body plan

Nevertheless, despite the effective adaptive morphological attributes discussed above, *L. helicolumna* retains the characteristic leptomitid body plan. The family Leptomitidae is exclusively composed of thin-walled, single spongocoel species, in which the skeleton is mainly constructed by long longitudinal monaxons (Figure 3). Some authors have proposed that they possessed an asconoid aquiferous system^39,41^, although this remains difficult to verify. However, if our interpretation is correct, the ability of *L. helicolumna* to close its osculum suggests that it functioned as a single-module organism, according to the definition of Fry^60^. According to the latter, an aquiferous module is an osculum and its associated sponge cells and aquiferous ducting^60^, and a true osculum is an exhalant pore whose opening and closing can be regulated by exopinacocytes alone^60^. This concept of aquiferous modules has been proved by experimental studies in living multi-oscular species^66^.

A modular organization endows sessile organisms with many benefits, such as unlimited growth, the plasticity of form, and a higher possibility of regeneration after a catastrophic impact^67,68^. Although there are single-modular sponges surviving today (e.g., *Sycon*), the modular body plan is far more prevalent^60,69^. A statistical analysis revealed that during the ∼20- Myr evolutionary span of archaeocyaths, an extinct reef-building poriferan group, there was a notable rise in both the prevalence of modular species and the complexity of their modularity^67^. The simple non-modular aquiferous system, such as that of *L. helicolumna*, was likely an intrinsic disadvantage when competing with modular species.

It is worth noting that whether the single spongocoel in other leptomitid taxa represents a single module remains uncertain. In the absence of evidence for a contractable osculum, the single spongocoel could alternatively represent a common exhalant canal shared by multiple modules^60^.

Phenotypic plasticity is another critical adaptive strategy for sessile organisms that cannot migrate when the environmental factors change^21,70,71^. The ecological advantage of plasticity in sponges can be generally manifested by the fact that demosponges, widely known for their morphological plasticity, constitute over 80% of the living sponge species^5^. Many demosponges are able to reorganize their shape, aquiferous system, and skeletal production to optimize their survival chances and energy costs in changing environmental conditions^19,72,73^. Even asconoid calcareans, such as *Leucosolenia* and *Clathrina*, are flexible in their growth form in terms of number and coalescence of the branching tubes^60^. In contrast, despite over a century of study^54,74^, leptomitids have consistently been described as thin-walled, solitary fossils with a regular shape and a single spongocoel, lacking evidence of branching or budding. Notably, the morphometric data in this study show that *L. helicolumna* maintains a fixed shape across different body sizes, exhibiting a strict isometric growth mode. It seems clear that *L. helicolumna* lacks the phenotypic plasticity that is ubiquitously seen in extant sponges. This may, in turn, negatively affect the survival of this species.

In some studies, the regular morphology in early Palaeozoic sponges has been interpreted as evidence of more complex genetic regulation, while the seemingly undetermined morphology in living sponges was viewed as a result of secondary simplification^41,75^. Indeed, being morphologically flexible is highly advantageous for sponge survival, and the molecular mechanisms regulating the morphological plasticity are likely complex. They involve the integration of multiple modules, the sensing of the changing environmental factors, the timing of the morphological reorganization, and a trade-off between nutrient and energy costs^19,67^.

## CONCLUSION

The newly described early Cambrian leptomitid *Lotispongia helicolumna* gen. et sp. nov. exhibits several sophisticated functional morphological innovations that appear well adapted to the shallow-water siliciclastic environment. Key features – such as rope-like longitudinal spicule bundles, a thick root tuft, radiating spicules, and a contractable osculum – are previously unrecorded in other leptomitids, yet many are analogous to strategies adopted by extant sponges. Despite these novelties, *L. helicolumna* retains a characteristic leptomitid body plan: a strictly single-spongocoel (indicative of a single-modular organization in the species) and the lack of phenotypic plasticity. These features contrast with the modularity and morphological plasticity widely recognized as ecologically advantageous in modern sessile organisms. This case study highlights both the evolutionary conservatism of the thin-walled, regular, and single- spongocoel body plan of leptomitids and the potential for substantial adaptive innovations within this ancient body plan.

## LIMITATIONS OF THE STUDY

This study analyzed the pros and cons in the functional morphophology of *Lotispongia helicolumna* gen. et sp. nov. and argued that the lack of modularity and phenotypic plasticity may be drawbacks of its body plan. This hypothesis is to be tested in the future when statistic data of fossil spatio-temporal distribution accumulates.

## RESOURCE AVAILABILITY

All data have been presented in the main text and supplemental materials. All investigated fossils are now deposited at the Nanjing Institute of Geology and Palaeontology.

## ACKNOWLEDGEMENTS

We appreciate the constructive suggestions and comments from Joachim Reitner, Christine Schönberg, Huilong Ou, and the three anonymous reviewers of an earlier version of this manuscript submitted to Proceedings B. We thank Jean-Bernard Caron for allowing access to the fossil specimen illustrated in Figure S1. This study was supported National Natural Science Foundation of China (42330209, 41972016).

## AUTHOR CONTRIBUTIONS

C. L. collected specimens, performed data analysis, and wrote this manuscript; Y. H. performed data analyses; Z. S. collected specimens and studied the trilobite biostratigraphy; H. S. collected specimens and studied the hyolith biostratigraphy; S.-C. L. provided data on the living sponge; T. W. and L. Z. investigated the spicule illustrated in Figure S1; F. Z. provided the donated specimens. All authors read and improved the manuscript.

## DECLARATION OF INTERESTS

The authors declare no competing interests.

## MATERIALS AND METHODS

The material sources and fossil locality have been introduced in “Preservation of the sponge fossils”. Here is some supplementary information on their biostratigraphy. The sponge fossils co-occur with the trilobites *Megapalaeolenus deprati* Mansuy^76^ (Figures 1C and 1D), *Redlichia yunnanensis* Resser & Endo^77^ (Figure 1I), and *Breviredlichia granulosa* Zhang & Lin in Yin & Li^78^ (Figure 1J). Among them, *M. deprati* and *B. granulosa* typically correspond to the upper part of the Wulongqing Formation, while *R. yunnanensis* to the lower part^32,79^. The overlap of these species at the sponge fossil interval indicates that the studied materials belong to the middle part of the Wulongqing Formation. In addition, *B. granulosa* is correlative to the *Megapalaeolenus fengyangensis* Zone in the middle Yangtze shelf^80,81^, indicating an age of early Cambrian Age 4^31,82^ (Figure 1B). Consistent with the trilobite biostratigraphy, the co-occurring hyolith “*Linevitus*” *guizhouensis* (Figures 1E and 1F) has been described from the coeval Balang Formation of Guizhou Province, and *Doliutheca orientalis* (Figures 1G and 1H) from the Shipai Formation of the eastern Yangtze Gorges^83,84^.

All fossils illustrated here were investigated using a Nikon SMZ18 microscope coupled with a Nikon DSRi2 camera and photographed using a Nikon D850 single-lens reflex camera. The morphometric measurements were conducted based on macroscopic and microscope photos using ImageJ.

The sponge spicule illustrated in Figure S2 was macerated from the black shale of the Shuijingtuo Formation (Cambrian Stage 3), from the locality reported in Luo et al.^18^ It was then embedded in resin, polished to exposed the cross-section, and studied using a Zeiss Crossbeam 550 Scanning Electronic Microscope coupled with an Oxford UltimM170 Energy-Dispersive X-ray Spectroscopy. The elemental distribution was mapped at a voltage of 15 kV.

## SUPPLEMENTAL INFORMATION

**Figure S1** Preservation of *Pirania muricata* (ROM 56249), where an open-based spicule was reported^41^. (A) Overview of the specimen. (B, C) Closer views of the open-baed spicule. This object is preserved in a black mineral, less compacted, and shows transverse fractures. In contrast, the *Pirania* spicules are strongly flattened, covered by a reflective film (probably clay minerals), and do not show transvers fractures. (D) Another black fragment on the same bedding plane, showing similar preservation as the brachiopod and the open-based spicule. (E) Another part of *P. muricata*, where spicules are also preserved with strong flattening and associated with a reflective film. (F) The open-based spicule is photographed under cross-polarized light.

**Figure S2** Cross-section of a well-preserved spicule from the black shale of the Shuijingtuo Formation (Cambrian Stage 3), the same locality as the fossils reported in Luo et al.^18^(A) Overview of the cross-section. (B–F) A closer view (B) and its elemental maps. The spicule was basically composed of silica (C–F), and the concentric structure is formed by differentiated solution of silica (B–C) and the filling of carbon in the soluted pores (E). The carbon signal outside of the spicule comes from the resin in which the spicule was embedded.

**Table S1** List of investigated fossils, with raw data of morphometric measurements.

**Table S2** Selected specimens of *Lotispongia helicolumna* used for the morphometric analysis.

## Notes

### Competing Interest Statement

The authors have declared no competing interest.

