## Supplemental Figures S1+S2 for "Advanced adaptive strategies in an ancestral body plan: insights from a 510-Ma-old leptomitid sponge"

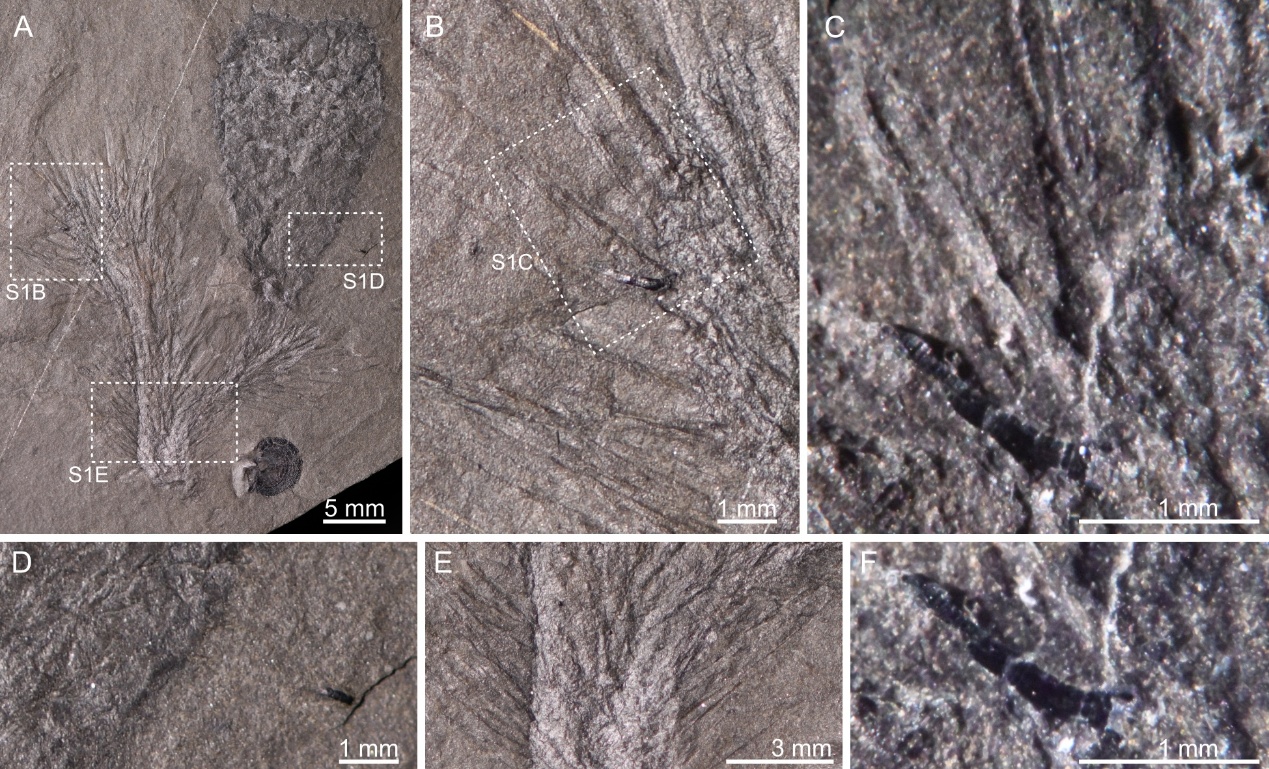

**Figure S1** Preservation of *Pirania muricata* (ROM 56249), where an open-based spicule was reported^41^. (A) Overview of the specimen. (B, C) Closer views of the open-baed spicule. This object is preserved in a black mineral, less compacted, and shows transverse fractures. In contrast, the *Pirania* spicules are strongly flattened, covered by a reflective film (probably clay minerals), and do not show transvers fractures. (D) Another black fragment on the same bedding plane, showing similar preservation as the brachiopod and the open-based spicule. (E) Another part of *P. muricata*, where spicules are also preserved with strong flattening and associated with a reflective film. (F) The open-based spicule is photographed under cross-polarized light.

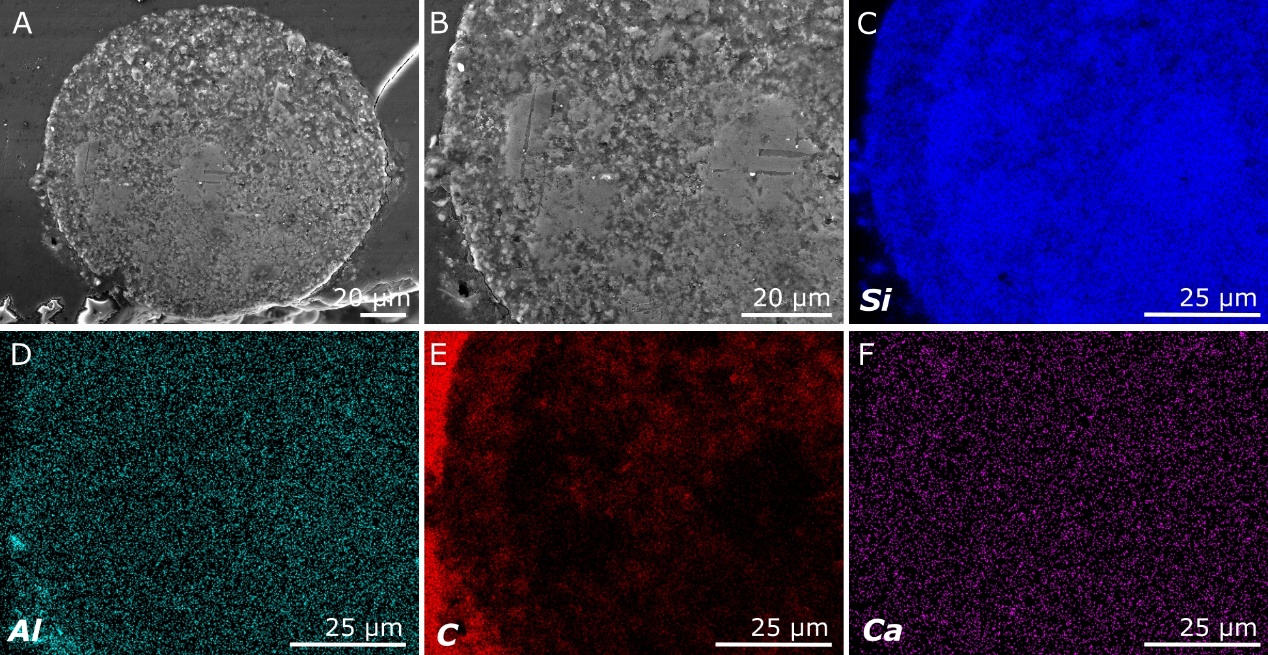

**Figure S2** Cross-section of a well-preserved spicule from the black shale of the Shuijingtuo Formation (Cambrian Stage 3), the same locality as the fossils reported in Luo et al.^18^(A) Overview of the cross-section. (B–F) A closer view (B) and its elemental maps. The spicule was basically composed of silica (C–F), and the concentric structure is formed by differentiated solution of silica (B–C) and the filling of carbon in the soluted pores (E). The carbon signal outside of the spicule comes from the resin in which the spicule was embedded.
